# GacA regulates symbiosis and mediates lifestyle transitions in *Pseudomonas*

**DOI:** 10.1101/2025.01.20.633850

**Authors:** Youqing Luo, Apsara Srinivas, Casey Guidry, Carolee Bull, Cara H. Haney, Corri Hamilton

**Author notes:** Co-first authors contributed equally to publication; author order was determined alphabetically (Youqing Luo, Apsara Srinivas).

## Abstract

Through horizontal gene transfer, closely related bacterial strains have assimilated distinct sets of genes, resulting in significantly varied lifestyles. However, it is not clear how strains are able to properly regulate horizontally transferred virulence genes. We hypothesized that strains may use components of the core genome to regulate diverse horizontally acquired genes. To investigate how closely related bacteria assimilate and activate horizontally acquired DNA, we used a model of *Pseudomonas fluorescens* subspecies *brassicacearum* strains, N2E2 and N2C3, which exhibit contrasting lifestyles on the model plant Arabidopsis. *P. brassicacearum* strain N2E2 is a plant commensal and contains the genes that encode biosynthetic enzymes for the anti-fungal compound 2,4-Diacetylphloroglucinol (DAPG). In contrast, N2C3 lacks DAPG biosynthesis and has gained a pathogenic island encoding syringomycin (SYR)-and syringopeptin (SYP)-like toxins from the plant pathogen *P. syringae.* This causes a transition in lifestyle from plant-protective N2E2 to plant pathogenic N2C3. We found that N2E2 and N2C3 share a highly conserved two-component system GacA/S, a known regulator of DAPG and SYR/SYP. Using knockout mutations, we found that a *ΔgacA* mutation resulted in loss of expression of SYR/SYP virulence genes and returned pathogenic N2C3 to a plant commensal lifestyle. Our study further explored the conservation of regulatory control across strains by demonstrating that GacA genes from both distant and closely related *Pseudomonas* strains could functionally complement one another across the genus.

**IMPORTANCE:** Emerging pathogens represent a significant threat to humans, agriculture, and natural ecosystems. Bacterial horizontal gene transfer aids in the acquisition of novel genes that facilitate bacterial adaptation to new environments. Our work shows a novel role for GacA in orchestrating the regulatory changes necessary for virulence and lifestyle transitions facilitated by HGT. These findings suggest that the GacA/S system plays a key role in mediating transitions across diverse *Pseudomonas* symbiotic lifestyles. It provides insights into the mechanisms that drive the emergence of pathogenic strains and highlights potential targets for managing bacterial threats to plant health.

## INTRODUCTION

Horizontal gene transfer (HGT) is a powerful evolutionary mechanism that enables bacteria to rapidly acquire virulence traits and adapt to new hosts, presenting a major threat as new pathogenic strains emerge (1–3). Unlike vertical inheritance, HGT allows bacteria to integrate genetic material directly from other species or environmental sources, leading to dramatic shifts in bacterial behavior, including the acquisition of genes associated with antibiotic resistance, metabolic versatility, and virulence (2). This rapid adaptability makes HGT a significant driver of emerging pathogenic threats, as formerly benign or beneficial strains can quickly develop mechanisms to harm their hosts by gaining pathogenic capabilities.

The *Pseudomonas fluorescens* in the *P. brassicacearum* subclade (*P. brassicacearum* hereafter), along with species in the closely related *P. corrugata* and *P. mediterranea* subclades, exemplifies this threat of pathogen emergence from commensal lifestyle. Strains within these subclades share 99% 16S rRNA identity and range from beneficial to pathogenic lifestyles in association with diverse hosts. The *P. brassicacearum* subclade includes WCS365, NFM421, DF41, and N2E2 which are commensal strains that promote root growth and protect plants from diverse fungal and bacterial pathogen infections (4–8). However, the *P. corrugata* subclade includes pathogenic strains including *P. corrugata,* the causative agent of tomato pith necrosis (9) and strains within the *P. brassicacearum* subclade, including N2C3 and R401, cause root stunting on gnotobiotic Arabidopsis (10, 11). Plant pathogenic strains in both *P. corrugata* and *P. brassicacearum* use a lipopeptide toxin-based virulence strategy that results in disease on diverse hosts ranging from plant species from the families *Brassicaceae* and *Papaveroideae* (9, 10). Studies on *P. brassicacearum* N2C3 show that it acquired the toxin-encoding pathogenicity islands through HGT (10). This clade illustrates how HGT facilitates the sudden emergence of virulent pathogens capable of threatening diverse ecosystems.

One central challenge in understanding the rise of new pathogens is uncovering how bacteria regulate these newly acquired genes to ensure they function cohesively with existing cellular processes. While the mechanics of HGT are well understood, an open question remains: how are these foreign genes effectively regulated to ensure functionality within their new host? Bacteria rely on two-component systems (TCS), like the GacS/GacA system, to sense and respond to environmental signals, modulating gene expression accordingly (12). In *Pseudomonas* spp., the GacS/GacA system regulates hundreds of genes, through the Rsm family of small trans-acting regulatory RNAs, including those tied to plant pathogenicity and biocontrol (12, 13). In beneficial strains like *P. protegens* CHAO and Pf-5 GacA regulates antifungal secondary metabolites such as diacetylphloroglucinol (DAPG) (14, 15), while in pathogens like *Pseudomonas syringae* pv. *syringae*, GacA modulates virulence traits including the type III secretion system (16, 17). This led us to hypothesize that GacA/S may regulate newly acquired virulence genes in both pathogenic and commensal strains to facilitate lifestyle transitions between closely related strains.

In this study, we aimed to understand how closely related strains within *P. brassicacearum* utilize a functional GacA/S system to specialize on hosts and regulate either pathogenic or commensal traits acquired through horizontal gene transfer. By examining the gain and loss of *gacA* through homologous recombination, we investigated how these genetic changes drive transitions between pathogenesis and commensalism. We used genetic complementation across distantly related *Pseudomonas* strains and characterized natural mutations to assess GacA’s role in these lifestyle shifts. By focusing on closely related strains in a subclade with both commensals and pathogens, our study highlights the importance of the GacA/S system in regulating host specialization and the manifestation of either beneficial or harmful traits. Through advancing our understanding of these regulatory mechanisms, we aim to inform strategies for managing emerging pathogens and protecting plant health.

## RESULTS

### GacA regulates recently horizontally acquired pathogenicity and specialized metabolite genes in *Pseudomonas brassicacearum*

To better understand how otherwise commensal strains can correctly utilize pathogenic genomic elements gained through horizontal gene transfer, we focused on characterizing the commensal *P. fluorescens* strains in the *P. brassicacearum* subclade, namely two closely related isolates from groundwater, plant pathogenic (N2C3) and plant commensal (N2E2) (10, 18). Phylogenetic analysis indicates that these strains are so closely related that they would share the same Operational Taxonomic Unit (OTU) based on 16S rRNA amplicon sequencing as the well-characterized antifungal and plant growth-promoting strain WCS365 (10). We found that while GacA/S system is conserved across the genus *Pseudomonas*, downstream mobile elements including DAPG, and the SYR/SYP toxins, show rapid gain and loss between closely related strains [(10), Fig. 1A].

**Figure 1:**
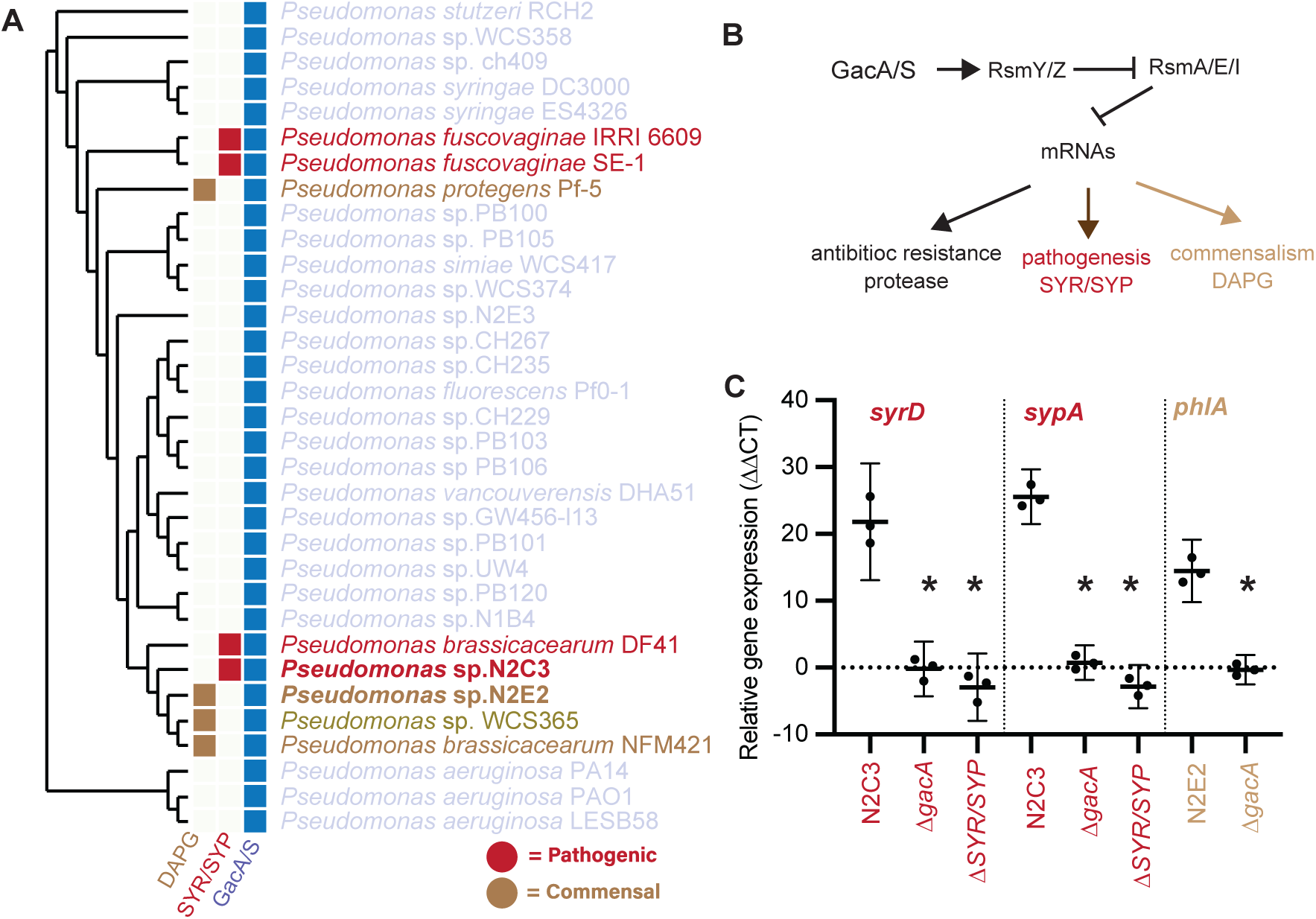
The presence of a functional GacA/S system is necessary for expression of horizontally transferred genomic islands in *P. brassicacearum*. A) A phylogenetic tree based on 122 core *Pseudomonas* genes shows conservation of GacA/S while DAPG and Pathogenicity Island gained and lost throughout the genus B) Model of the hypothesis of the role of Gac-Rsm is regulation of horizontally acquired genomic elements. C) ΔΔCT values of genes *syrD* (syringomycin), *sypA* (syringopeptin), *phlA* (2,4-diacetylphloroglucinol), and compared to the *recA* housekeeping gene in Kings B Media. Each dot represents an average of three technical replicates and one biological replication (n = 9). Shown are means +/− standard deviation and asterisk for significance (*p* < 0.05) in a Student’s t-test.

To determine whether *gacA/S* regulates horizontally acquired islands in the *P. brassicacearum,* including SYR/SYP (Fig. 1B), we generated deletions of *gacA* in N2E2 and N2C3 and tested whether loss of *gacA* affects the expression of genes encoding the biosynthetic enzymes that produce DAPG, SYR/SYP, and the associated anti-fungal and plant virulence phenotypes. Using qRT-PCR, we found that deletion of *gacA* in *P. brassicacearum* N2C3 decreases expression of *syrD* and *sypA,* required for biosynthesis of SYR and SYP respectively (Fig. 1C). Deletion of *gacA* in N2E2 resulted in decreased *phlA* required for biosynthesis of DAPG (Fig. 1C). These results confirm that *gacA* is required for expression of horizontally transferred symbiosis factors that regulate both beneficial and pathogenic traits with *P. brassciacearum*.

We then tested whether a loss of *gacA* would also affect the virulence and antifungal activities of N2C3 and N2E2. Consistent with its role in regulating the expression of SYR/SYP, we found that an N2C3 *ΔgacA* mutant loses virulence as measured by root stunting and reduced plant fresh weight (Fig. 2A-C). Consistent with the absence of SYR/SYP and other plant virulence factors in the commensal strain N2E2, we found no difference in root length and fresh weight in an N2E2 Δ*gacA* mutant (Fig. 2A-C). Similarly, we found that while wildtype N2C3 and N2E2 inhibit the growth of the oomycete root-rot pathogen *Aphanomyces euteiches* AE-1, the *ΔgacA* mutants in both N2E2 and N2C3 lose the ability to inhibit *A. euteiches* AE-1 growth (Fig. 2D-E). Collectively, these findings suggest that GacA may facilitate the regulation of horizontally transferred islands, including relatively recently acquired SYR/SYP and DAPG genes, which have been recently gained and lost in *P. brassicacearum*.

**Figure. 2:**
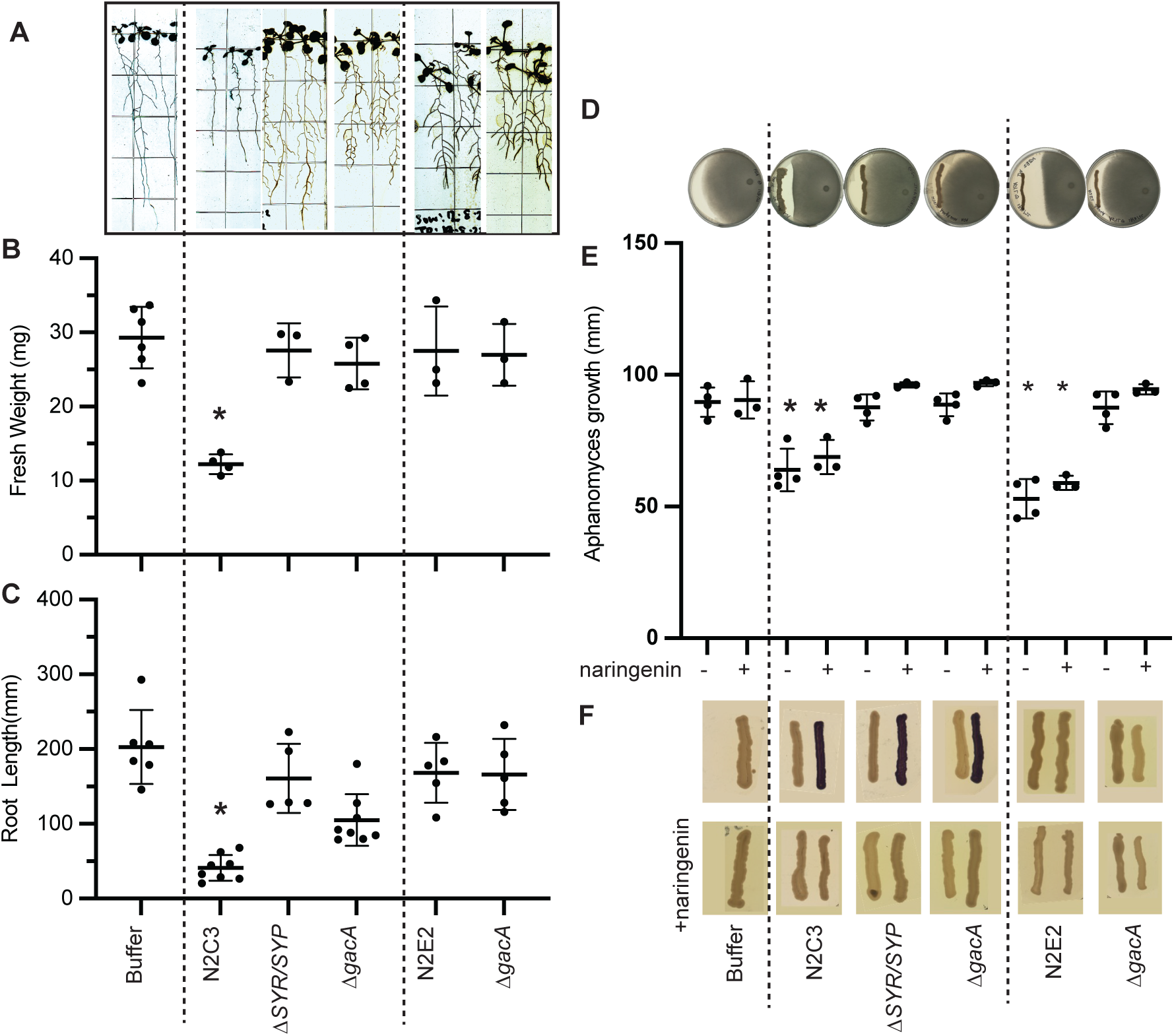
GacA controls lifestyle-dependent phenotypes independent of quorum signaling. A) Representative images of gnotobiotic Arabidopsis seedlings inoculated with Buffer or *P. brassicacearum* N2C3, N2C3 *ΔgacA*, N2C3 *Δsyr/syp,* N2E2, or N2E2 *ΔgacA*. B) Seedling root length and C) plant fresh weight was quantified. Each dot represents the average of 3-5 plants, and three to six biological replicates were performed for each treatment (n = 9-30). Mean +/− standard deviation is shown and asterisk indicates significant (p < 0.05) differences as determined by a one-way ANOVA followed by a post-hoc Tukey HSD test. D) Representative images of *Aphanomyces euteiches* AE-1 oomycete antagonism assay. E) Quantification of oomycete growth when competed with P. *brassicacearum* N2C3, N2C3 *ΔgacA*, N2C3 *Δsyr/syp,* N2E2, or N2E2 *ΔgacA* in the presence or absence of the quorum quencher naringenin. Experiments were repeated 4 independent times with 3 plates per replicate (n = 9-12). F) Representative images of qualitative quorum sensing activity detected by measuring the purple violacein pigment produced by *Chromobacterium violaceum* CV026 AHL biosensor reporter system in the presence (top) or absence (bottom) of naringenin.

SYR/SYP biosynthesis is regulated by a C6-acyl homoserine lactone (AHL) perceived by an associated LuxR regulator that is co-inherited with the SYR/SYP island (10). Some GacA/S-dependent processes are quorum dependent (19), suggesting that GacA/S may indirectly regulate SYR/SYP biosynthesis by regulating the associated *luxR/I* genes. We tested whether GacA regulates SYR/SYP and DAPG through quorum sensing by asking whether the N2C3 Δ*gacA* mutant still produces C6-AHL. We used *Chromobacterium violacium* CV026 C6-AHL reporter that turns purple in the presence of C6-AHLs; N2C3, but not N2E2, was previously shown to activate this reporter (10). We found that N2C3 *ΔgacA* mutant could still induce the reporter, indicating that it still makes C6-AHL (Fig. 3F). N2E2 in contrast, lacks production of C6-AHL and did not activate the reporter (Fig. 3F). To further investigate if suppression of quorum signaling would affect the production of SYR/SYP and DAPG and consequentially inhibition of oomycete pathogen growth, we used the quorum quenching flavanone naringenin, which reduces AHL levels (20). While naringenin suppressed N2C3-mediated activation of the CV026 reporter *in vitro* (Fig. 3F), we found that both N2C3 and N2E2 were able to inhibit *Aphanomyces* growth in the presence of naringenin (Fig. 3E). These data indicate that the production of DAPG and SYR/SYP secondary metabolites, and GacA regulation of SYR/SYP and DAPG, are independent of quorum signaling mechanisms (Fig. 3E, F).

**Figure. 3:**
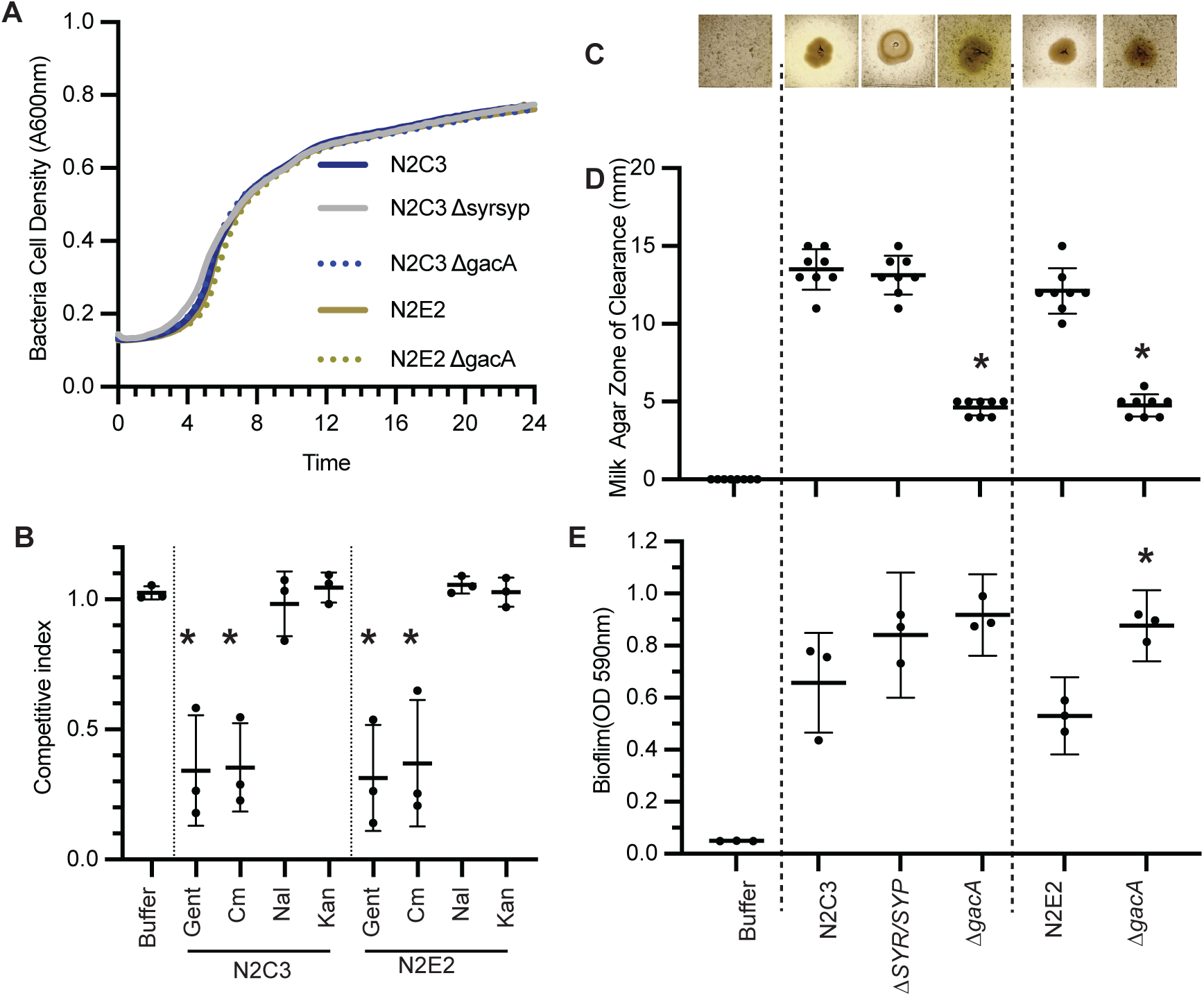
GacA controls the regulation of conserved symbiosis factors in closely related strains with divergent lifestyles. A) 24-hr growth curves of each strain were performed in LB media with no significant differences in growth over time via repeated measures of ANOVA (p < 0.05; n = 9). B) To test whether *gacA* is differentially required for antibiotic resistance we grew N2C3, N2C3 *ΔgacA*, N2E2, or N2E2 *ΔgacA* in subinhibitory concentrations of multiple antibiotics and described growth outcome as the competitive index of *ΔgacA* strains compared to the wildtype parent strain. C-D) To test whether *gacA* is necessary for protease activity, 5 µl of 0.02 OD_600nm_ bacterial cultures were spotted on milk agar plates. C) Representative images of N2C3 and controls on milk agar. D) Quantification of zone of clearance for *P. brassicacearum* N2C3, N2C3 *Δsyr/syp*, N2C3 *ΔgacA*, N2E2, or N2E2 *ΔgacA*. Each dot represents an average of 3 spots. E) To test whether *gacA* is necessary for biofilm formation, a PVC-crystal violet assay was performed and biofilm formation was quantified colorimetrically via absorbance 590. B,D-E) Experiments were repeated 3 to 6 independent times (n = 9-20). Mean shown and asterisk indicate significant (p < 0.05) differences as determined by a one-way ANOVA follow by a post-hoc Tukey HSD test.

### GacA functions in shared symbiotic traits independent of bacterial lifestyle

Because we found that GacA regulates lifestyle-dependent horizontally transferred islands, we hypothesized that it also might regulate a shared set of genes involved in symbiosis independent of lifestyle. Importantly, we found that mutations in *gacA* do not lead to overall growth defects in minimal media and rich media (Fig. 3a and Fig. S1). GacA is known to regulate antibiotic resistance in pathogenic *Pseudomonas* stains (21). We found that even within *P. brassicacearum* strains, loss of *gacA* increases sensitivity to subinhibitory concentrations of gentamycin and chloramphenicol (Fig. 3B and Table S1). Protease activity and biofilm formation are important for symbiosis; biofilm formation is required for plant colonization and for virulence (22), while protease activity is required for rhizosphere competition and utilized to lyse a portion of the cell walls of pathogenic fungi (23). To quantify protease activity, we used a milk agar assay and found that GacA is necessary for protease activity in both *P. brassicacearum* N2C3 and N2E2 (Fig. 3C-D). Using a crystal violet assay, we found that the loss of *gacA* significantly increases biofilm formation in N2E2 and trends toward being increased in N2C3 (*p* < .0932) (Fig. 3E). These data suggest that GacA plays a role in shared functions useful for symbiosis but is not required to maintain overall growth.

### Complementation of GacA between strains restores symbiotic functions

To determine if a single GacA protein can regulate both pathogenic and beneficial islands we asked whether *gacA* from the N2C3 pathogenic strain could complement an N2E2 *ΔgacA* commensal mutant and vice-versa. We generated constructs where *gacA* was expressed under its native promoter on a plasmid. As expected, *N2C3-gacA_pro_:N2C3-gacA* complemented the *in planta* N2C3 *ΔgacA* phenotype to wildtype-like levels for root length, fresh weight, and protease activity (Fig. 4A-C). Likewise, *N2E2-gacA_pro_:N2E2-gacA* complemented the protease activates of the N2E2 *ΔgacA* mutant back to N2E2 wildtype levels (Fig. 4C). We found that introduction of the beneficial *N2E2-gacA_pro_:N2E2-gacA* construct into the N2C3 *ΔgacA* mutant also complimented protease activity to the same level as *N2C3-gacA_pro_:N2C3-gacA* construct (Fig. 4A-C). This held true for the introduction of *N2C3-gacA_pro_:N2C3-gacA* into the N2E2 *ΔgacA* mutant, which restored the protease activity to N2E2 wildtype levels (Fig. 4A-C). The ability of GacA to regulate lifestyle-specific traits in diverse strains underscores its importance in mediating transitions between different lifestyles.

**Figure 4.**
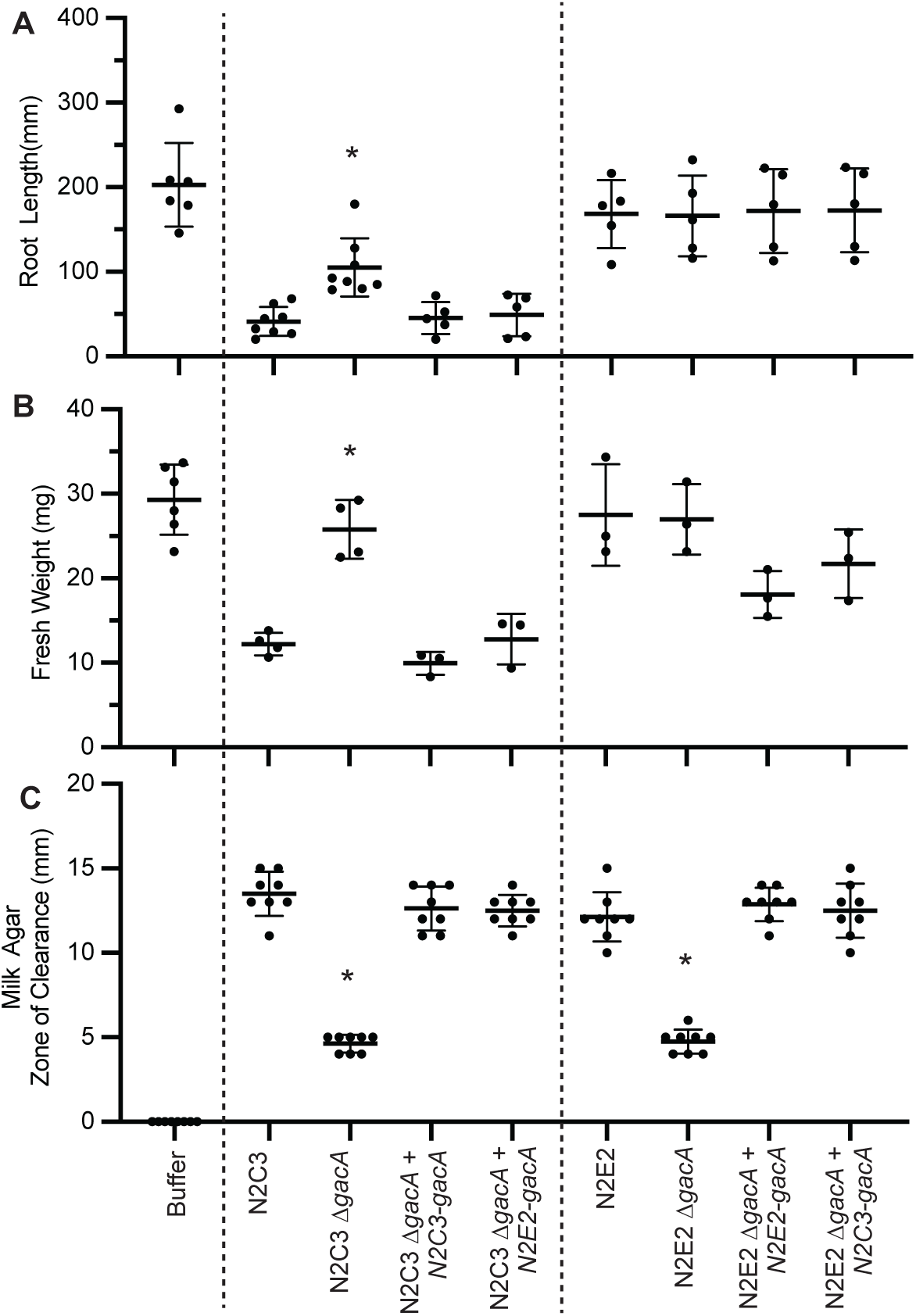
GacA can heterologously complement *Pseudomonas* with distinct lifestyles. To test whether virulence phenotypes in N2C3 can be complemented by *gacA* from N2E2, gnotobiotic Arabidopsis seedlings were inoculated with P. *brassicacearum* N2C3, N2C3 *ΔgacA*, express *gacA* under its native promoter *N2C3-gacA_pro_:N2C3-gacA* or *N2E2-gacA_pro_:N2E2-gacA*. Differences were quantified using A) seedling root length and B) plant fresh weight. Each dot represents average of 3-5 plants over three to six biological replicates (n = 9-30). Mean +/− standard deviation is shown and asterisks indicate significant (*p* < 0.05) differences as determined by a one-way ANOVA follow by a post-hoc Tukey HSD test. C) To test whether *gacA* is sufficient for protease activity, 5 µl of 0.02 OD_600nm_ cultures were spotted on milk agar plates, and the zone of clearance was measured. Mean +/− standard deviation is shown and asterisks indicate significant (*p* < 0.05) differences as determined by a one-way ANOVA followed by a post-hoc Tukey HSD test.

### GacA can heterologously regulate the expression of virulence factors across *Pseudomonas* species

To determine if GacA from N2C3 or N2E2 can regulate virulence in distantly related *Pseudomonas* spp., we tested whether *P. brassicacearum gacA* could heterologously regulate gene expression in a syringomycin-producing *P. syringae* strain 485-10, originally isolated from citrus blast infected tissues (24). We found the *P. syringae* 485-10 showed protease activity on milk agar indicating it has a functional GacA/S signaling pathway (Fig. 5A). We found that like syringomycin-producing N2C3 and *P. fuscovaginae* SE-1, that *P. syringae* 485-10 caused reduced fresh weight when inoculated onto Arabidopsis seedlings (Fig. 5B).

**Figure 5.**
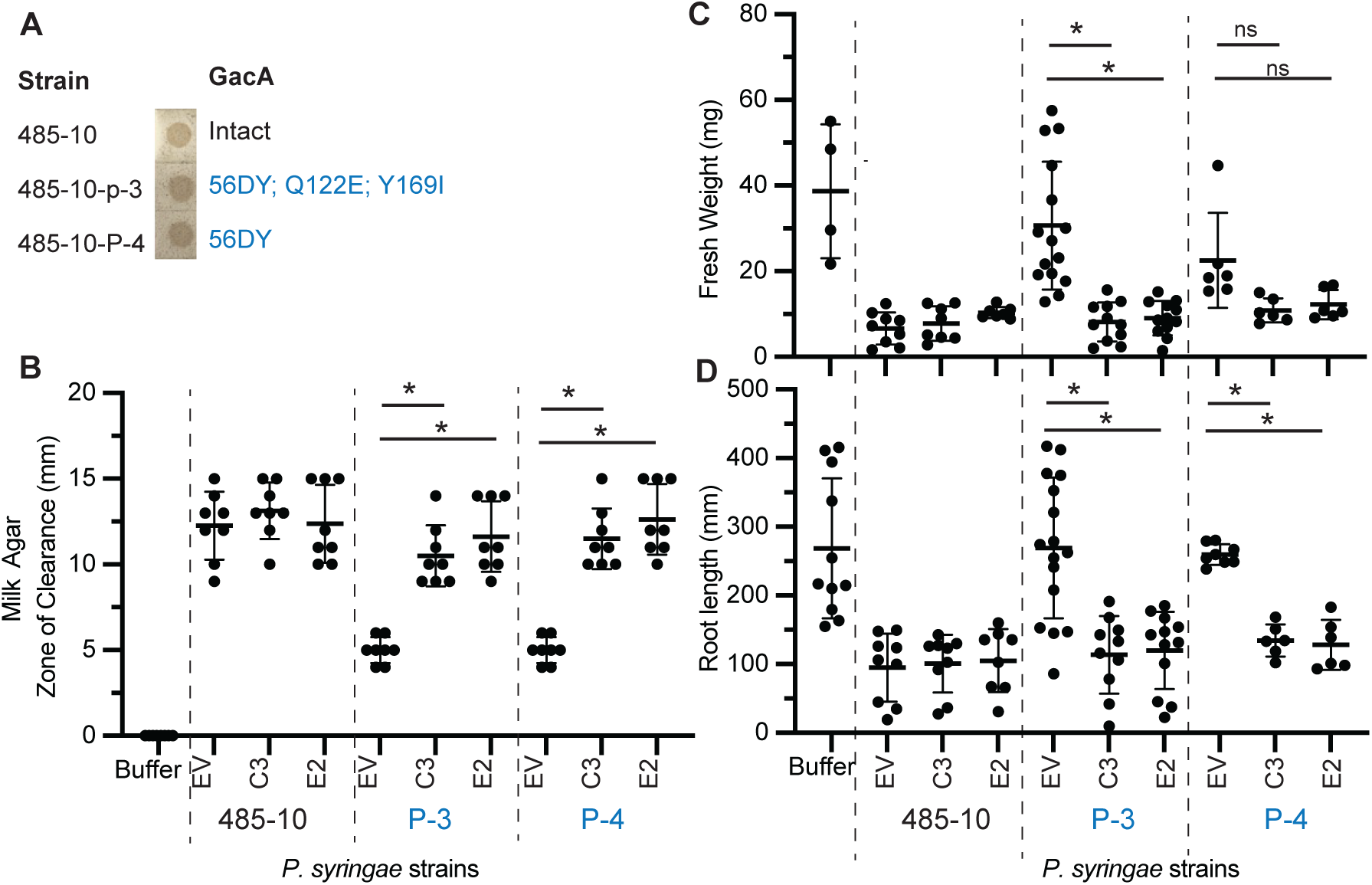
*gacA* from *P. brassicacearum* can heterologously complement *P. syringae* 485-10 *gacA* mutants. A) Spontaneous protease-negative *P. syringae* strains mutants were identified in *P. syringae* 485-10. The *gacA* genotype is indicated by Sanger sequencing. B) Protease activity, C) plant fresh weight and D) root growth inhabitation of 485-10 with introduction of empty vector (EV), *N2C3-gacA_pro_:N2C3-gacA* (C3) or *N2E2-gacA_pro_:N2E2-gacA* (E2). Mean +/− standard deviation is shown and asterisks indicate significant (*p* < 0.05) differences as determined by a one-way ANOVA followed by a post-hoc Tukey HSD test.

To identify GacA mutants, two spontaneous protease activity-negative *P. syringae* 485-10 mutants were identified by screening on milk agar (Table S2). Spontaneous protease-negative mutants 485-10-P-3, and 485-10-P-4 lost the ability to clear milk agar resulting in a reduced zone of clearance compared to respective parent strains (Fig. 5A, C). By PCR amplification and Sanger sequencing we found that spontaneous protease mutants 485-10-P-3 and 485-10-P-4 had missense mutations (Fig. 5A). Consistent with a loss of GacA/S signaling, when inoculated onto plants, all protease-negative mutants resulted in higher fresh weight compared to its respective parent strains indicating a loss of virulence (Fig. 5B).

To determine if a *P. brassicacearum gacA* gene could regulate virulence traits in *P. syringae,* we tested whether N2C3 and N2E2 *gacA* genes can complement a *P. syringae gacA* mutant. We found that in *P. syringae* 485-10-P-3 and 485-10-P-4 *gacA* mutants that *N2C3-GacA* or *N2E2-GacA* restored the fresh weight, root length and protease activity (Fig. 5B-D). These data indicate that restoration of GacA in distantly related *Pseudomonas* complements *in vitro* and *in planta* phenotypes. This restoration of pathogenic traits in GacA-deficient strains illustrates the role of GacA in facilitating the acquisition of virulence traits across the genus *Pseudomonas*.

## DISCUSSION

The GacS/GacA two-component system (TCS) plays a crucial role in determining the lifestyle of *Pseudomonas* species, regulating the delicate balance between pathogenicity and commensalism. This study focuses on the *Pseudomonas brassicacearum* subclade, particularly the closely related strains N2E2 and N2C3, to uncover the specific role of GacA in managing newly acquired HGT-driven traits that facilitate rapid shifts in behavior that can transform previously harmless strains into potent pathogens.

Our results reveal that GacA is central to the activation of virulence factors, particularly those acquired through horizontal gene transfer. For instance, in the pathogenic strain N2C3, GacA controls the expression of syringomycin and syringopeptin—key secondary metabolites within a pathogenic island transferred from other *Pseudomonas* species (Fig. 2). Disruption of GacA in N2C3 abolishes these pathogenic traits, effectively reverting the strain to a commensal state (Fig. 2A-C). This aligns with existing literature, which has demonstrated GacA’s role in regulating virulence across various *Pseudomonas* species, including *Pseudomonas syringae*. It is known that spontaneous mutants occur in rich media (Bull et al. 2001). In general, soils are nutrient deprived. Importantly, when a nutrient rich environment like a root is present, bacteria may be more likely to toggle and decrease the numbers of cells that are virulent? This would indicate that syringomycin plays a different role in soil bacteria than in foliar pathogens. It may protect the bacterium from some organisms or inhibit fungi to remove competition in nutrient poor conditions. Investigating the varied environmental triggers for this mutation may provide insight into the role of the regulated metabolites in fitness of the organism in the varied environments they occupy.

The study also highlights GacA’s broader regulatory influence, extending beyond specific pathogenic traits to include general symbiotic functions that are vital for bacterial survival and interaction with plant hosts. This includes the regulation of protease activity, biofilm formation, and the production of antifungal compounds like 2,4-diacetylphloroglucinol (DAPG). In N2E2, for example, the loss of GacA results in reduced expression of DAPG and a subsequent decrease in fungal antagonism (Fig. 2D-F), demonstrating GacA’s role in maintaining the beneficial aspects of bacterial-plant interactions. This suggests that GacA’s regulation of antifungal activity is crucial for enhancing plant defense mechanisms, thereby strengthening the symbiotic relationship between bacteria and their plant hosts. DAPG has been shown to be vital for the biocontrol efficacy of *P. fluorescens* in suppressing soil-borne pathogens like *Fusarium* species. Fungal antagonism plays a crucial role in bacterial symbiosis with plants by helping bacteria suppress pathogenic fungi, thereby promoting plant health and facilitating beneficial plant-microbe interactions.

Our heterologous complementation experiments further underscore the versatility and importance of GacA across different strains. When GacA from either N2E2 or N2C3 was reintroduced into gacA-deficient mutants, the mutants regained their lost phenotypic traits, such as root length, fresh weight, and protease activity. Notably, this complementation was effective even when GacA was expressed under non-native promoters, indicating that the regulatory mechanisms governed by GacA are robust and conserved across different strains. This finding is particularly relevant in the context of horizontal gene transfer, where newly acquired genes require a regulatory framework like GacA to be effectively integrated into the bacterium’s existing genetic and functional landscape.

Moreover, the ability of GacA to restore symbiotic functions in distantly related *Pseudomonas syringae* highlights its broad regulatory influence. In our study, 4 spontaneous protease activity-negative mutants were complemented with GacA vectors from N2E2 or N2C3, leading to the partial or complete restoration of their original phenotypes. These data indicate that GacA from *P. brassicacearum* can regulate virulence components in distantly related species, suggesting *P. brassicacearum* is also capable of expressing virulence factors from distantly related strains. As it is likely that SYR/SYP in *P. brassicacearum* N2C3 was horizontally acquired from *P. syringae* (10), these findings suggest that a functional GacA/S pathway in *P. brassicacearum* might facilitate these types of HGT events.

The implications of this study extend beyond the specific strains examined, offering insights into the broader mechanisms of bacterial evolution and adaptation. In agricultural contexts, where *Pseudomonas* species are integral to both plant health and disease, understanding and manipulating GacA function could provide new strategies for enhancing beneficial plant-microbe interactions or mitigating pathogenic threats. The GacS/GacA TCS, through its regulation of small non-coding RNAs (sRNAs) like RsmY and RsmZ, controls the expression of genes associated with virulence, secondary metabolite production, and stress responses. Our work highlights GacA’s role as a key regulatory component that supports bacterial adaptation in diverse environments, bridging symbiotic and pathogenic lifestyles. In doing so suggest further research into this pathway, especially in relation to different bacterial lifestyles, could lead to the development of next-generation microbial inoculants designed to promote plant growth and suppress disease, ultimately contributing to more sustainable agricultural practices.

## MATERIALS AND METHODS

### Bacteria growth conditions and selection conditions

All *Pseudomonas* strains were cultured at 28℃ using Lysogeny Broth (LB) or agar plates. When appropriate, LB was supplemented with 10% sucrose, 25 µg/ml kanamycin (Km), 30 µg/ml gentamicin (Gm), and 10 µg/ml nalidixic acid.

### Growth curve analysis

*P. brassicacearum* wild-type strains, respective *ΔgacA*, and complementation strains were inoculated in LB (24) in a standard flat-bottom 96-well plate at 28 °C with a starting 0.02 OD 600_nm_ culture in three technical replicates. The bacterial growth was measured spectrophotometrically at 600_nm_ in a continuously shaking plate.

### Strain construction

*E. coli* DH5α was used for the construction and maintenance of plasmids, and *E. coli* SM10 λpir was used for biparental conjugation with *P. brassicacearum*. Deletion mutants were generated using the two-step allelic exchange method as previously described (25, 26). Deletion constructs for N2E2 Δ*gacA* and N2C3Δ*gacA* were generated using suitable primers (Table S3).

Genomic DNA was extracted using the Qiagen Puregene kit A. Regions 500-1000 bp upstream and downstream of the target gene were amplified with High-Fidelity Phusion Polymerase (NEB) and purified using the Qiagen QIAquick® PCR Purification Kit. These regions were connected via one-step overlap PCR and digested with the appropriate restriction enzymes (Table S3). The PCR products were then ligated into the pEXG2 suicide vector containing *sacB* and transformed into *E. coli* SM10 λpir Transformants were selected on LB agar with 30 µg/ml Gm, and the plasmid was transferred to *Pseudomonas* strains via conjugation, integrating into the genome through homologous recombination.

The *gacA* complementation plasmids were created using the pBBR1MCS-2 plasmid as a backbone (27) The N2E2 *gacA* and N2C3 *gacA* genes, along with approximately 200 bp upstream of each gene, were amplified using specific primers (Table S3). These constructs were then electroporated into *Pseudomonas* strains to generate the *gacA* complementation strains (28). Kanamycin (Km) was used for selection and maintenance of the plasmid. Wild-type and *ΔgacA* strains with an empty vector (EV) were generated similarly by electroporating pBBR1MCS-2 into electrocompetent cells. These cells were prepared by pelleting, washing, and resuspending cultures in 300 mM sucrose. Transformants were selected on and maintained using Km (25 µg/ml) for *P. brassicacearum*. *Pseudomonas syringae* cultures were grown in LB medium at 28°C with shaking (200 rpm). After an initial 2-day growth, 1 mL was transferred to 50 mL of fresh ½-strength LB and grown overnight. This culture was used to inoculate fresh ½-strength LB (50 mL) and grown to an OD_600_ of 0.5–0.6. Exponential-phase cells were washed with 0.5 M sucrose, resuspended in 1 mL of 0.5 M sucrose, and mixed with 5 µg plasmid DNA (100 µL). Electroporation was performed at 25 µF, 200 Ω, and 9.5 kV cm⁻¹ with a ∼5 ms time constant. Cells were incubated in LB for 2 h, and transformants were selected on LB plates containing 25 µg/mL kanamycin after 2 days of growth at 28°C (29).

### *In planta* vertical plate disease assays

*Arabidopsis thaliana* Col-0 seeds were sterilized by gas sterilization for 1.5 hrs. 7 mL of 37% HCl was added to 150 mL of 6% bleach to generate Cl2 gas and sterilized seeds were stored in 0.1% sterile phytoagar at 4 °C in the dark for four days before sowing. ½ Murashige and Skoog (MS) medium plates with 1x MES buffer and 1% sucrose were used to germinate seeds. Plates were then placed vertically in a 23°C growth room with 16 hrs light, 8 hrs dark. Five days post germination, seedlings were then transplanted to no-sucrose ½ x MS plates with 1% MES buffer. Roots were inoculated after five days of transplanting. All the bacteria were diluted (0.001 OD_600nm_) before inoculation (10). The fresh weight of inoculated seedlings was measured six days after inoculation.

### In vitro assays

#### Antagonism assay (30)

*Aphanomyces euteiches* AE-1 was maintained on Potato Dextrose Agar (PDA) plates, with sub-culturing done by transferring a 5 mm agar plug from an actively growing colony to fresh PDA plates, which were incubated at 25°C for 7-10 days. Bacterial strains were cultured on LB plates and incubated at 28°C for 24 hours. Single colonies were scrape inoculated onto antagonism PDA plate. Plates were incubated at 25°C for 7-10 days, and the zone of inhibition around each well was measured to assess the antagonistic effect of the bacterial strains against *A. euteiches*.

#### Biofilm assays (31)

Overnight culture of *P. brasscicacearum* N2C3 or N2E2 wildtype or *ΔgacA*, mutant cells were diluted (0.1 OD 600_nm_) in M63 medium (1× M63 salt, 0.2% glucose, 0.5% casamino acids, and 1 mM MgSO4) and 100 μL of diluted cultures were incubated at 27 °C for 18 h in non-tissue culture-treated 96-well plates. Subsequently, the plate was rinsed, stained with 125 μL of 0.1% crystal violet for 10 min, and dried overnight. The crystal violet was dissolved in 125 μL of 30% acetic acid for 10 min and transferred to a new 96-well, flat bottom plate. Absorbance was measured at 550 nm using a spectrophotometer (SpectroMax) and all absorbance signals were averaged and normalized against the wildtype values.

#### Proteases activity assay (31)

5 µL of a bacterial suspension was spotted onto the center of each milk agar plate. Plates were incubated at 28°C for 72 hours (32). Zones of clearance around the bacterial colonies were measured, indicating the proteolytic activity of the bacteria. Milk agar plates were prepared by adding 20% (v/v) sterile skim milk to autoclaved nutrient agar at approximately 50°C, then pouring the mixture into sterile Petri dishes. Plates were allowed to solidify at room temperature.

##### Quorum Sensing Assay

Bacteria were cultured overnight in LB broth with the appropriate antibiotics at 28°C with shaking at 200 rpm. The reporter strain culture was spread onto plates and allowed to dry. Subsequently, 10 µL of the adjusted signal-producing strain culture and control autoinducer were spotted onto the agar plates. The plates were incubated at 28°C for 48 hours. The presence and intensity of purple pigment indicates the level of quorum sensing signal induction (10, 33). Quorum quenching media was supplemented with 0.1% (w/v) naringenin (Sigma-Aldrich) (20).

#### Gene expression analysis

Transcripts of genes were quantified via RT-qPCR as described previously (34). RNA was extracted from Kings B liquid cultures using the Qiagen RNAeasy kit. cDNA was synthesized from normalized RNA using Promega reverse transcriptase. qPCR was performed in triplicate with at least 3 biological replicates, and gene expression levels were normalized using the ΔΔCt method with *recA* as the housekeeping gene

### Sequencing

To confirm the presence and integrity of the *gacA* gene, we performed Sanger sequencing. Genomic DNA was extracted from bacterial cultures using the Qiagen DNeasy Blood & Tissue Kit. The gacA gene was amplified using high-fidelity Phusion Polymerase (New England Biolabs) and specific primers flanking the *gacA* coding region (Supplementary Dataset S1). Purified PCR products were submitted to Azenta for Sanger sequencing. Sequences were to reference sequences in the NCBI database using BLAST to confirm the identity and integrity of the *gacA* gene.

### Statistics

Statistical analyses were conducted to evaluate differences in bacterial growth and disease symptoms across various experimental treatments. All statistical tests were performed using GraphPad Prism software. The type of analysis applied depended on the experimental design. Student’s t-test was used for pairwise comparisons to identify significant differences between two groups. Analysis of Variance (ANOVA) was employed for experiments involving multiple groups. For data collected across multiple time points or under repeated measures, a repeated measures ANOVA was used to account for the correlation within subjects over time. Post-hoc tests were applied when ANOVA results indicated significance to identify specific group differences. For all tests, statistical significance was defined as p<0.05. Detailed descriptions of the statistical tests used, including their assumptions and the conditions under which they were applied, are provided in the text where results are presented. Results are reported with appropriate measures of central tendency and variability (e.g., mean ± standard deviation of the mean). Graphical representations of the data include error bars to reflect this variability, ensuring clarity and reproducibility in the interpretation of findings.

## ACKNOWLEDGEMENTS

This work was supported by CIHR Grant (PJT -169051) and NRCan (SPP-144-1) grants awarded to C.H.H. and was supported by the USDA National Institute of Food and Agriculture Federal Appropriations under project #PEN04926 (accession 7006350) awarded to C.T.B.

